# TSLP-mediated inflammatory factors production contributes to metastatic breast cancer-induced bone cancer pain in mice

**DOI:** 10.1101/423673

**Authors:** Ying Chen, Pei-Feng Xian, Sheng-Xu Wang, Lu Yang

## Abstract

**Objective:** Metastatic breast cancer-induced bone cancer pain (BCP) is one of the most disabling factors in patients suffering. Recent studies show thymic stromal lymphopoietin (TSLP) is a proinflammatory factor and high expression in cancer cells. Here we investigated whether and how TSLP released form breast cancer cells contributes to BCP.

**Methods:** Braset cancer cells MDA-MB-231 were intramedullary injected into the femur to induce BCP, TSLP neutralizing antibody was intrathecally injected and pain hypersensitivity was checked by behavioral testing. The TSLP expression was knockdown by shRNA. Human TSLP, TNF-α, IL-1β, IL-6, CCL2, CXCL2 and CXCL5 in intramedullary space of the femur of rats was detected by ELISA.

**Results:** TSLP neutralizing antibody intrathecally injection significantly reduced MDA-MB-231 cells inoculation induced persistent mechanical allodynia and heat hyperalgesia and TSLP increased. Furthermore, Persistent mechanical allodynia and heat hyperalgesia of mice models were also decreased in MDA-MB-231 cells with TSLP knockdown. Finally, some inflammatory factors, TNF-α, IL-1β, IL-6, CCL2, CXCL2 and CXCL5 were significantly elevated in the intramedullary space lavage fluid of the right femur after cancer cell inoculation, but were obviously reduced in models with neutralizing antibody of TSLP treatment and TSLP knockdown.

**Conclusion:** TSLP-mediated inflammatory factors release contributes to the maintenance of tumoral hypersensitivity. Inhibition of the TSLP may provide a new therapy for BCP management.

## Introduction

Epidemiological study found that bone metastasis was a common complication of breast cancer patients[1]. After metastasized into bone, nociceptors innervating of bone are activated by cancer cells and some factors like nerve growth factor (NGF)[2], prostaglandin E2 (PGE2)[3] and endothelin[4] which secreted by cancer cells, as well as tumor associated immune cells, osteoblasts, osteoclasts lead bone cancer pain production. But current treatment strategies often can’t provide adequate analgesia and cause unacceptable side effects[5]. So further understanding the underlying mechanisms of the development of BCP is important for effectively treating these patients

Accumulating evidence suggests that inflammatory factors, like TNF-α, IL-1β and so on, have an important role in the induction and maintenance of chronic cancer pain[6-8]. After nerve injury or proinflammatory mediators’ stimulation, spinal microglia is activated and products a large number of inflammatory factors, and that leads to chronic cancer pain[9]. Thymic stromal lymphopoietin (TSLP) is a master switch for allergic inflammation and promotes primarily type 2 inflammatory responses that as a characteristic of a large number of inflammatory factors production, such as TNF-α and IL-1β[10-12]. Recently studies found that TSLP could activate neurons to induce itch[13] and had a high expression in cancer cells[14, 15]. However, whether TSLP play a role in BCP is unknown.

In this study, we used neutralizing antibody of TSLP to treat MDA-MB-231 cell inoculation-induced BCP model and examined TSLP levels in the intramedullary space of the right femur of model. We also constructed BCP model used MDA-MB-231 cell with the TSLP expression knockdown and examined inflammatory factors levels in the intramedullary space of the right femur of this model.

## Methods

### Cell culture and generation of stable cell lines

Normal breast epithelium MCF-10A and breast carcinoma cell lines MCF-7, MDA-MB-231, T47D, BT-549, ZR-75-30 and BT-474 were all purchased from ATCC (Manassas, VA). Briefly, these cell lines were cultured in Dulbecco’s modified Eagle’s medium or RPMI 1640 media supplemented with 10% fetal bovine serum (Life Technologies/Invitrogen, South America) in 5% CO_2_ at 37°C. MDA-MB-231 cells were infected with a GPU6/GFP/Neo-shTSLP vector and selected by Neomycin (1000 ng/mL) to generate polyclonal cells with stable knockdown of TSLP. shRNA-vectors targeting TSLP were purchased from GenePharma (Shanghai, China) and transfected into cells using Lipofectamine 2000 (Invitrogen, USA)[16].

### Animals and tumor inoculation

Adult (8 weeks) females C57Bl/6 mice were used to construct BCP model according to the guidelines of the International Association for the Study of Pain and approved by the Animal Care and Use Committee of Southern Medical University. All mice had free access to food and water with a 12/12 light/dark cycle. Breast cancer cell lines, MDA-MB-231, was purchased from the Cell Bank of Type Culture Collection of Chinese Academy of Sciences (Shanghai, China). The MDA-MB-231 cells were collected and resuspended in phosphate buffered saline (PBS) in a concentration of 5× 10^7^ cells/mL. The animals were anesthetized with sodium pentobarbital (50 mg/kg, i.p.). Anarthrotomy was done to expose the condyles of the distalfemur. PBS containing 10^6^ RM-1 cells (20 μl) was injected into the intramedullary space of the right femur with a 30 G needle and the injection site was sealed with bone wax. Sham control mice were injected with same amount of PBS[17].

### Behavioral analysis

Behavioral analysis was performed as described previously[17, 18].

### ELISA

20μl 0.9% physiological saline was injected in intramedullary space of the right femur of rats with a 30 G needle and about 15μl lavage fluid was collected for ELISA. Human TSLP, TNF-α, IL-1β, IL-6, CCL2, CXCL2 and CXCL5 ELISA kit was purchased from R&D. ELISA was performed according to manufacturer’s protocol. The standard curve was included in each experiment.

### Western blot

Western blot studies were performed as described previously[16, 19]. The antibodies were shown as follows: TSLP antibody (1:1000, rabbit, Abcam, USA) and β-actin antibody (1:1000, mouse Proteintech, USA). IRDye800 conjugated anti-rabbit IgG and IRDye680 conjugated anti-mouse IgG (LiCor, USA) were used as secondary antibodies. Signal intensities were analyzed by using the Odyssey infrared Image System (LiCor, USA). The densitometry results were first normalized with that of β-actin and then compared with the control to obtain relative fold changes.

### Statistical analysis

Statistical analysis was performed as described previously[17]. A value of *P*<0.05 was considered to indicate a statistically significant difference in all statistical analyses.

## Result

### Neutralizing antibody of TSLP attenuates MDA-MB-231 cell inoculation-induced pain hypersensitivity

To investigate the role of endogenous TSLP in BCP, MDA-MB-231 cells were inoculated into the intramedullary space of mice femur to construct a mice model of BCP and checked pain behaviors. Pain behavioral studies showed that tumor cell inoculation produced an obvious pain hypersensitivity compared with PBS inoculation (Sham group), which was characterized by heat hyperalgesia (the paw withdrawal latency, PWL) (fig. 1A) and mechanical allodynia (the paw withdrawal threshold, PWT) (fig. 1B) in the right hindpaws of inoculated mice. But, when the TSLP neutralizing antibody was intrathecally injected to mice model of BCP at 7 days after inoculation, the PWL and PWT were partly attenuated, and maintained to day 28 (fig. 1A and B). Meantime, we found that the TSLP levels was not significant change between day 0 and day 28 in Sham animals, but significantly increased at day 28 of MDA-MB-231 cells inoculation which were regulated with intrathecally injection of TSLP neutralizing antibody (fig. 1C). These data suggest TSLP is involved in tumor cell inoculation-induced pain hypersensitivity.

**Fig. 1.**
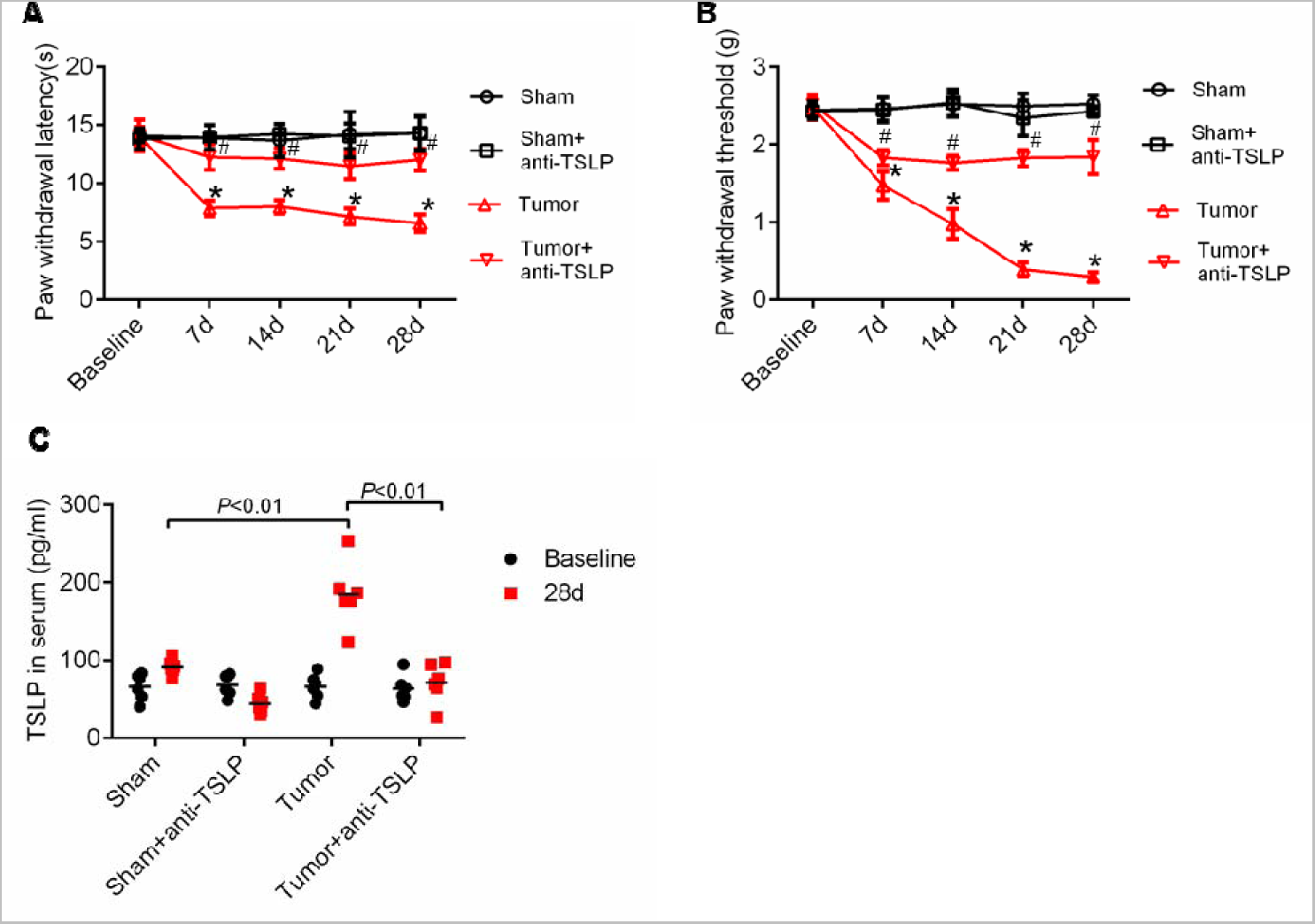
Neutralizing antibody of TSLP attenuates MDA-MB-231 cell inoculation-induced pain hypersensitivity. The PWL (A) and PWT (B) of behavioral tests show that mice displayed both heat hyperalgesia and mechanical allodynia in the ipsilateral paw after MDA-MB-231 cell inoculation. (C) The TSLP levels in the intramedullary space of the right femur of mice. ^∗^*P* <0.01, *vs*. Sham group. #*P* <0.01 *vs*. Tumor group. n=6 mice per group.

### MDA-MB-231 cell inoculation induced pain hypersensitivity by releasing TSLP

To check whether the increased TSLP of the intramedullary space of the right femur of mice was released by breast cancer cells, the TSLP protein of normal breast epithelium (MCF-10A) and breast cancer cell lines (BT-474, T-470, MCF-7, ZR-75-30, BT549 and MDA-MB-231) was examined. As shown in fig.2A, we found that the expression of TSLP protein was higher levels in breast cancer cell lines than that in MCF-10A. On the line with that, the TSLP secretion of breast cancer cell lines also was more than MCF-10A (fig.2B). Moreover, when MDA-MB-231 cells with TSLP knockdown were inoculation to mics, the TSLP levels of the intramedullary space lavage fluid were obviously decreased (fig.2C) and MDA-MB-231 cell inoculation induced pain hypersensitivity was significantly alleviated (fig.2C and D). These data indicated that the higher levels of TSLP released by breast cancer cells induced pain hypersensitivity.

**Fig. 2.**
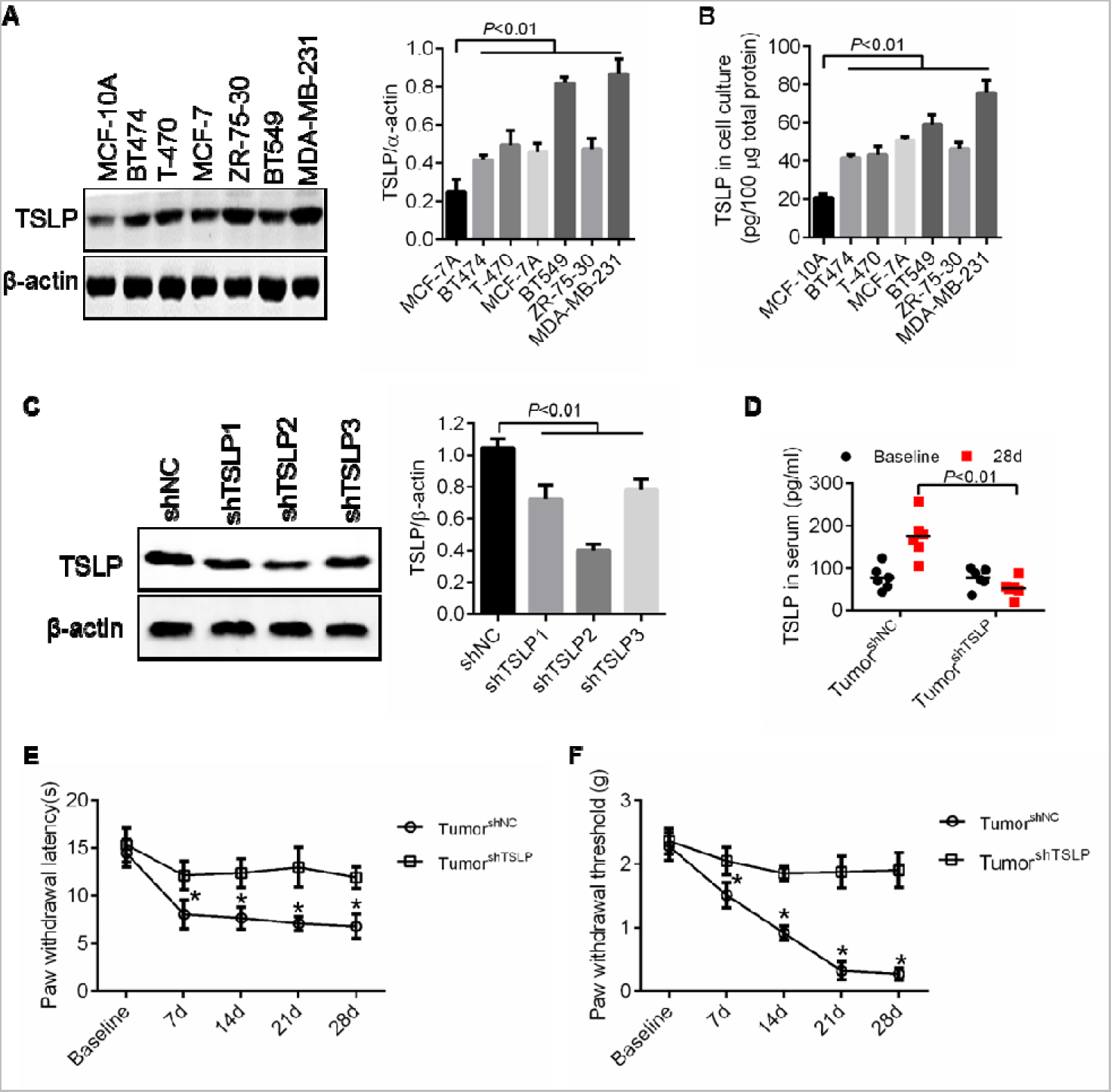
MDA-MB-231 cell inoculation induced pain hypersensitivity by releasing TSLP. (A) The expression of TSLP protein in breast cancer cell lines. (B) Elisa was performed to estimate the TSLP levels in the supernatant of cell culture medium. (C) The expression of TSLP were knockdown by plasmid with shTSLP. (D) The TSLP levels in the intramedullary space lavage fluid of the right femur of rats BCP model. The PWL (E) and PWT (F) of behavioral tests show that mice displayed both heat hyperalgesia and mechanical allodynia in the ipsilateral paw after MDA-MB-231 cells inoculation. ^∗^*P* <0.01, vs. Tumor^shNC^ group. n = 6 mice per group.

### Inhibition of TSLP decreased neuroinflammatory factors levels

Accumulating evidence suggests that neuroinflammation has an important role in the induction and maintenance of chronic pain[6] and TSLP is a proinflammatory factor which induced TH2 inflammation[20]. So we further investigated whether TSLP induced and maintained pain hypersensitivity by promoting neuroinflammatory factors production. Elisa shown that the levels of TNF-α, IL-1β, IL-6, CCL2, CXCL2 and CXCL5 were significantly increased in the intramedullary space lavage fluid of the right femur of rats with MDA-MB-231 cells inoculation for 28d (fig. 3 A-F). But treatment with Neutralizing antibody of TSLP inhibited the increased levels of TNF-α, IL-1β, IL-6, CCL2, CXCL2 and CXCL5 compared with MDA-MB-231 cells inoculation that without treatment. Moreover, inoculation with MDA-MB-231 cells that TSLP was knockdown, the increased levels of TNF-α, IL-1β, IL-6, CCL2, CXCL2 and CXCL5 were significantly suppressed. These data illustrated that TSLP induced chronic pain by promoting neuroinflammatory factors levels production.

**Fig. 3.**
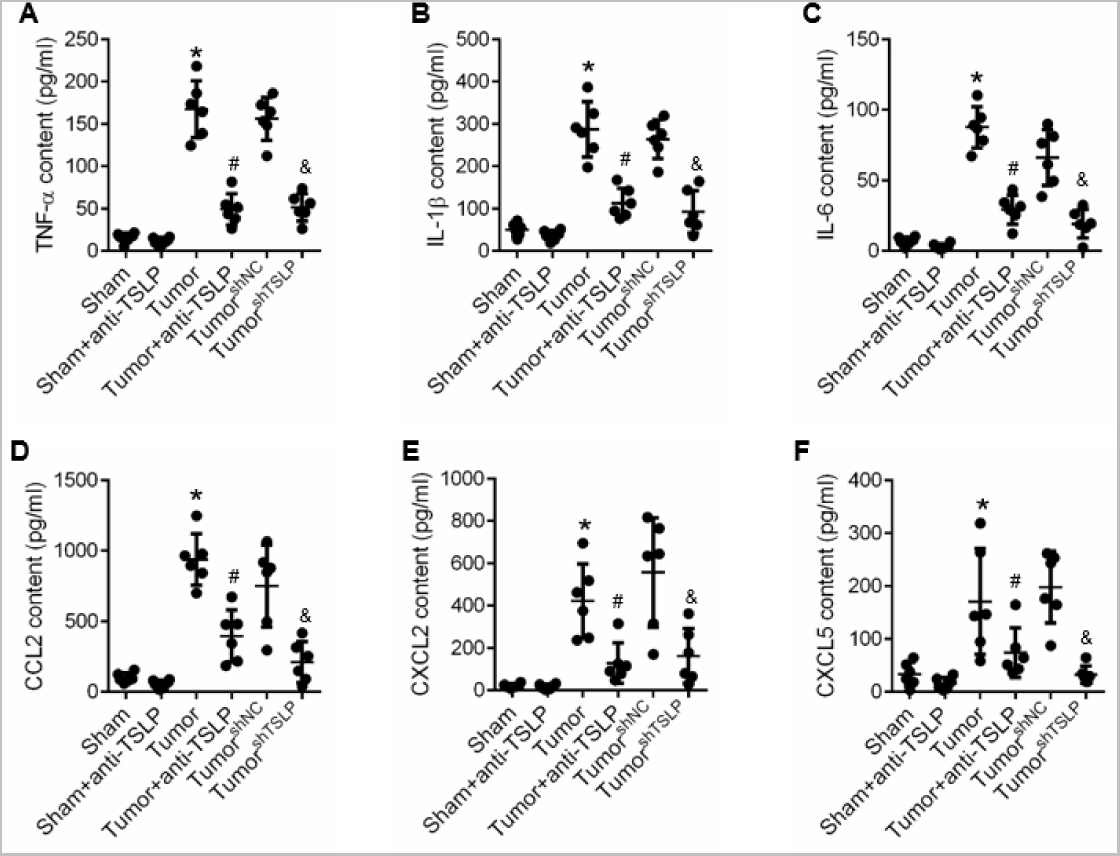
Neuroinflammatory factors in the intramedullary space lavage fluid of the right femur of rats BCP model. Elisa was performed to estimate the (A) TNF-α, (B) IL-1β, (C) IL-6, (D) CCL2, (E) CXCL2 and (F) CXCL5 levels in the intramedullary space lavage fluid of the right femur of rats BCP model after MDA-MB-231 cell inoculation for 28d. ^∗^*P* <0.01, *vs*. 28d Sham group. #*P* <0.01 *vs*. 28d Tumor group. &*P* <0.01 *vs*. 28d Tumor^shNC^ group. n = 6 mice per group.

## Discussion

The data presented herein demonstrate that TSLP was significant increase in the intramedullary space lavage fluid of the right femur after cancer cell inoculation. Inhibition of TSLP or cancer cell with TSLP knockdown attenuated inoculation-induced pain hypersensitivity. Our data also shows that some inflammatory factors, TNF-α, IL-1β, IL-6, CCL2, CXCL2 and CXCL5, which could induction and maintenance of chronic cancer pain, were significantly elevated in the intramedullary space lavage fluid of the right femur after cancer cell inoculation, but were obviously reduced in models with neutralizing antibody of TSLP treatment and TSLP knockdown. These data suggest that TSLP which secreted by cancer cell plays an important role in the induction and maintenance of BCP via promoting production of inflammatory factors.

TSLP is an important cytokine to regulate Th2 inflammation[20] and induce an abundant inflammatory factors secretion. Recent research findings that TSLP acts directly on a subset of TRPA1-positive sensory neurons to trigger robust itch behaviors in atopic dermatitis[13]. In our studies, we also found that inhibition of TSLP attenuated inoculation-induced pain hypersensitivity. These data indicated that TSLP play an important role in sensation of peripheral nervous.

However, what secret the increased TSLP in the intramedullary space lavage fluid of the right femur after cancer cell inoculation. Recent studies have also implicated cancer tissue TSLP expression and serum TSLP level were significantly increased[15, 21]. Moreover, the role of TSLP in tumor growth and metastasis is supported by many studies[14, 22, 23]. Consistent with these studies, we found that breast cancer cell lines have more expression and secretion than normal breast epithelium. In addition, TSLP content was also significant increase in the intramedullary space lavage fluid of the right femur after cancer cell inoculation, but the TSLP content was reduced in cancer cell with TSLP knockdown. All these suggested that breast cancer cells released an abundant TSLP to the intramedullary space lavage fluid of the right femur after cancer cell inoculation.

It is increasingly recognized that inflammatory mediators in the peripheral and central nervous system have a key role in the induction and maintenance of chronic pain[6]. Nociceptor neurons express the receptors include GPCRs, ionotropic receptors, and tyrosine kinase receptors, for all these inflammatory mediators, which act on their respective receptors on peripheral nociceptor nerve fibers and causes hypersensitivity and hyperexcitability of nociceptor neurons[24, 25]. We also found that many inflammatory mediators, like TNF-α, IL-1β, IL-6, CCL2, CXCL2 and CXCL5 were significantly increased in space lavage fluid of tumor models, but were obviously reduced in models with neutralizing antibody of TSLP treatment and TSLP knockdown. These data suggested that TSLP may promote inflammatory mediators’ production to induce BCP.

In this study, we found that inoculation of MDA-MB-231 cells into mouse femur induced bone pain hypersensitivity and chemokine TSLP was increased in the intramedullary space lavage fluid of femur. In association with these changes, Inhibition of TSLP or cancer cell with TSLP knockdown attenuated inoculation-induced pain hypersensitivity. Moreover, some increased inflammatory factors, which could induction and maintenance of chronic cancer pain, were significantly reduced in models with neutralizing antibody of TSLP treatment and TSLP knockdown. Our data suggest that TSLP-mediated inflammatory factors release contributes to the maintenance of tumoral hypersensitivity. TSLP may serve as a novel target for the treatment of metastatic breast cancer-induced BCP.

**Competing Interests** The authors declare that there is no conflict of interests.

## References

[1] Coleman R E. Clinical features of metastatic bone disease and risk of skeletal morbidity[J]. Clin Cancer Res. 2006, 12(20 Pt 2): 6243s-6249s.

[2] Miyagi M, Ishikawa T, Kamoda H, Suzuki M, Inoue G, Sakuma Y, Oikawa Y, Uchida K, Suzuki T, Takahashi K, Takaso M, Ohtori S. The efficacy of nerve growth factor antibody in a mouse model of neuropathic cancer pain[J]. Exp Anim. 2016, 65(4): 337-343.

[3] Frauenfelder S R, Freiberger S N, Bouwes B J, Quint K D, Genders R, Serra A L, Hofbauer G F. Prostaglandin E2, Tumor Necrosis Factor alpha, and Pro-opiomelanocortin Genes as Potential Mediators of Cancer Pain in Cutaneous Squamous Cell Carcinoma of Organ Transplant Recipients[J]. JAMA Dermatol. 2017, 153(3): 350-352.

[4] Smith T P, Haymond T, Smith S N, Sweitzer S M. Evidence for the endothelin system as an emerging therapeutic target for the treatment of chronic pain[J]. J Pain Res. 2014, 7: 531-545.

[5] Mantyh P. Bone cancer pain: causes, consequences, and therapeutic opportunities[J]. Pain. 2013, 154 Suppl 1: S54-S62.

[6] Ji R R, Xu Z Z, Gao Y J. Emerging targets in neuroinflammation-driven chronic pain[J]. Nat Rev Drug Discov. 2014, 13(7): 533-548.

[7] Stosser S, Schweizerhof M, Kuner R. Hematopoietic colony-stimulating factors: new players in tumor-nerve interactions[J]. J Mol Med (Berl). 2011, 89(4): 321-329.

[8] Jimenez-Andrade J M, Mantyh W G, Bloom A P, Ferng A S, Geffre C P, Mantyh P W. Bone cancer pain[J]. Ann N Y Acad Sci. 2010, 1198: 173-181.

[9] Ferrini F, Trang T, Mattioli T A, Laffray S, Del’Guidice T, Lorenzo L E, Castonguay A, Doyon N, Zhang W, Godin A G, Mohr D, Beggs S, Vandal K, Beaulieu J M, Cahill C M, Salter M W, De Koninck Y. Morphine hyperalgesia gated through microglia-mediated disruption of neuronal Cl(-) homeostasis[J]. Nat Neurosci. 2013, 16(2): 183-192.

[10] Wang Y H, Liu Y J. Thymic stromal lymphopoietin, OX40-ligand, and interleukin-25 in allergic responses[J]. Clin Exp Allergy. 2009, 39(6): 798-806.

[11] Ziegler S F, Artis D. Sensing the outside world: TSLP regulates barrier immunity[J]. Nat Immunol. 2010, 11(4): 289-293.

[12] Hammad H, Lambrecht B N. Barrier Epithelial Cells and the Control of Type 2 Immunity[J]. Immunity. 2015, 43(1): 29-40.

[13] Wilson S R, The L, Batia L M, Beattie K, Katibah G E, Mcclain S P, Pellegrino M, Estandian D M, Bautista D M. The epithelial cell-derived atopic dermatitis cytokine TSLP activates neurons to induce itch[J]. Cell. 2013, 155(2): 285-295.

[14] Olkhanud P B, Rochman Y, Bodogai M, Malchinkhuu E, Wejksza K, Xu M, Gress R E, Hesdorffer C, Leonard W J, Biragyn A. Thymic stromal lymphopoietin is a key mediator of breast cancer progression[J]. J Immunol. 2011, 186(10): 5656-5662.

[15] Watanabe J, Saito H, Miyatani K, Ikeguchi M, Umekita Y. TSLP Expression and High Serum TSLP Level Indicate a Poor Prognosis in Gastric Cancer Patients[J]. Yonago Acta Med. 2015, 58(3): 137-143.

[16] Yu C, Tang W, Wang Y, Shen Q, Wang B, Cai C, Meng X, Zou F. Downregulation of ACE2/Ang-(1-7)/Mas axis promotes breast cancer metastasis by enhancing store-operated calcium entry[J]. Cancer Lett. 2016, 376(2): 268-277.

[17] Xu J, Zhu M D, Zhang X, Tian H, Zhang J H, Wu X B, Gao Y J. NFkappaB-mediated CXCL1 production in spinal cord astrocytes contributes to the maintenance of bone cancer pain in mice[J]. J Neuroinflammation. 2014, 11: 38.

[18] Chaplan S R, Bach F W, Pogrel J W, Chung J M, Yaksh T L. Quantitative assessment of tactile allodynia in the rat paw[J]. J Neurosci Methods. 1994, 53(1): 55-63.

[19] Wang Y, Bu F, Royer C, Serres S, Larkin J R, Soto M S, Sibson N R, Salter V, Fritzsche F, Turnquist C, Koch S, Zak J, Zhong S, Wu G, Liang A, Olofsen P A, Moch H, Hancock D C, Downward J, Goldin R D, Zhao J, Tong X, Guo Y, Lu X. ASPP2 controls epithelial plasticity and inhibits metastasis through beta-catenin-dependent regulation of ZEB1[J]. Nat Cell Biol. 2014, 16(11): 1092-1104.

[20] Esnault S, Rosenthal L A, Wang D S, Malter J S. Thymic stromal lymphopoietin (TSLP) as a bridge between infection and atopy[J]. Int J Clin Exp Pathol. 2008, 1(4): 325-330.

[21] Ziegler S F, Roan F, Bell B D, Stoklasek T A, Kitajima M, Han H. The biology of thymic stromal lymphopoietin (TSLP)[J]. Adv Pharmacol. 2013, 66: 129-155.

[22] Vetter T, Borowski A, Wohlmann A, Ranjan N, Kuepper M, Badura S, Ottmann O G, Friedrich K. Blockade of thymic stromal lymphopoietin (TSLP) receptor inhibits TSLP-driven proliferation and signalling in lymphoblasts from a subset of B-precursor ALL patients[J]. Leuk Res. 2016, 40: 38-43.

[23] Barooei R, Mahmoudian R A, Abbaszadegan M R, Mansouri A, Gholamin M. Evaluation of thymic stromal lymphopoietin (TSLP) and its correlation with lymphatic metastasis in human gastric cancer[J]. Med Oncol. 2015, 32(8): 217.

[24] Gold M S, Levine J D, Correa A M. Modulation of TTX-R INa by PKC and PKA and their role in PGE2-induced sensitization of rat sensory neurons in vitro[J]. J Neurosci. 1998, 18(24): 10345-10355.

[25] Aley K O, Levine J D. Role of protein kinase A in the maintenance of inflammatory pain[J]. J Neurosci. 1999, 19(6): 2181-2186.

